## Appendix Table 1 for "Diet-driven differential response of *Akkermansia muciniphila* modulates pathogen susceptibility"

| **Appendix Table 1. Genome accession details for metatranscriptome analyses in Salmon.** | | | |
| --- | --- | --- | --- |
| **Organism Scientific Name** | **Organism Qualifier** | **Assembly Accession** | **Submission Date** |
| *Akkermansia muciniphila* | strain: ATCC BAA-835 | GCF_000020225.1 | 5/5/2008 |
| *Bacteroides caccae* | strain: ATCC 43185 | GCF_025146315.1 | 9/12/2022 |
| *Barnesiella intestinihominis* | strain: YIT 11860 | GCF_000296465.1 | 9/17/2012 |
| *Bacteroides ovatus* | strain: ATCC 8483 | GCF_001314995.1 | 10/15/2015 |
| *Bacteroides uniformis* | strain: ATCC 8492 | GCF_025147485.1 | 9/12/2022 |
| *Bacteroides thetaiotaomicron* | strain: DSM 2079 | GCF_014131755.1 | 8/10/2020 |
| *[Clostridium] symbiosum* | strain: ATCC 14940 | GCF_000466485.1 | 9/12/2013 |
| *Collinsella aerofaciens* | strain: JCM 10188 | GCF_010509075.1 | 2/13/2020 |
| *Faecalibacterium prausnitzii* | strain: A2165 | GCF_002734145.1 | 10/27/2017 |
| *Roseburia intestinalis* | strain: L1-82 | GCF_900537995.1 | 2/27/2019 |
| *Desulfovibrio piger* | isolate: FI11049 | GCF_900116045.1 | 11/12/2016 |
| *Marvinbryantia formatexigens* | strain: DSM 14469 | GCF_025148285.1 | 9/12/2022 |
| *Escherichia coli* | strain: HS | GCF_000017765.1 | 9/10/2007 |
| *[Eubacterium] rectale* | strain: VPI 0990 | GCA_022453685.1 | 3/3/2022 |
| *Citrobacter rodentium* | strain: ATCC 51459 | GCF_000835925.1 | 2/10/2015 |
